# Juxtaposition of human pluripotent stem cells with amnion-like cells is sufficient to trigger primitive streak formation

**DOI:** 10.1101/2025.07.11.664380

**Authors:** Xiangyu Kong, Anastasiia Nemashkalo, M. Cecilia Guerra, Miguel Angel Ortiz-Salazar, Elena Camacho-Aguilar, Aryeh Warmflash

**Affiliations:** Department of Biosciences, Rice University, Houston, TX 77005; Los Alamos National Laboratory, CINT/B11 Division, Los Alamos, NM; Department of Gene Regulation and Morphogenesis, Andalusian Center for Developmental Biology (CSIC-UPO-JA), Seville 41013, Spain; Department of Bioengineering, Rice University, Houston, TX 77005; Department of Developmental Biology, Washington University School of Medicine, St Louis, MO 63110; MRC Laboratory of Molecular Biology, Cambridge, UK

## Abstract

Studies in the mouse have established that communication between the trophectoderm and the epiblast is crucial for initiating gastrulation. In the primate embryo, the amnion rather than the trophectoderm is directly juxtaposed to the epiblast and may play this role. To model the interactions between the amnion and epiblast, we differentiated human pluripotent stem cells (hPSCs) to amnion-like cells (AMLCs) and juxtaposed them in a controlled manner with undifferentiated hPSCs, which represent the epiblast. We found that juxtaposition between these cell types is sufficient to initiate a range of cell behaviors associated with gastrulation including organized differentiation to primitive streak and downstream mesendodermal cell fates and directed cell migration out of the primitive streak region. Performing knockout experiments specifically in either the epiblast or amnion compartment revealed intricate crosstalk that is required to properly initiate gastrulation. In particularly, using knockouts of NODAL we show that Nodal signaling in both the amnion and epiblast is required for gastrulation patterning. Finally, we show that inductive ability is a transient property acquired during amnion differentiation, and that cells that differentiate from this inductive state acquire an extraembryonic mesenchyme identity. This study establishes a system to study epiblast-amnion communication and shows that this communication is sufficient to initiate gastrulation in the epiblast.

## Introduction

Gastrulation involves the differentiation of the pluripotent epiblast into the three germ layers of the embryo: ectoderm, mesoderm, and endoderm. In mammals, this germ layer differentiation occurs in the primitive streak, a structure which initiates in the posterior of the embryo at the boundary between embryonic and extraembryonic tissues and elongates anteriorly and distally. As cells transit through the primitive streak, they differentiate to mesendoderm and leave the epithelial epiblast layer, migrating in an anterior direction between the epiblast and the extraembryonic hypoblast.

Both in the mouse in vivo, and in human and mouse pluripotent stem cell (PSC) colonies in vitro, the events of gastrulation are initiated by a cascade of three signaling pathways: BMP, Wnt, and Nodal^1,2^. Loss of signaling through any of these pathways, for example, by deletion of the genes BMP4, Wnt3, or Nodal, causes a failure to gastrulate^3–5^. In mice, immediately prior to gastrulation, BMP4 is primarily expressed in the trophectoderm, and removal of the trophectoderm from the embryo proper also leads to a failure to express markers of gastrulation^6^. This has led to the model that gastrulation is triggered in the proximal posterior of the embryo by signals from the trophectoderm.

While mouse embryos are cup shaped with the epiblast and trophectoderm directly abutting and sharing a cavity prior to gastrulation, the embryos of primates, including humans, are disk shaped with a different relationship between the embryonic and extraembryonic tissues^7^. In primates, the amnion differentiates from the epiblast and forms the amniotic cavity prior to gastrulation^8^. Once present, the amnion forms a barrier between the epiblast and trophectoderm and might prevent signals from the trophectoderm from reaching the epiblast^9^. Thus, in primates, gastrulation-inducing signals such as BMP4 may initiate in the amnion rather than in the trophectoderm. Consistent with this idea, deletion of the amnion gene ISL1 in either non-human primate embryos or in a human pluripotent stem cell (hPSC) model of the amniotic sac reduces the expression of Bmp4 and impairs mesoderm formation and gastrulation^10^.

Recently, hPSC models based on micropatterning have allowed for dissection of the dynamics of signaling that lead to primitive streak induction and patterning at gastrulation stages^2,11–16^. In these models, cells are grown in colonies of defined size and shape and treated with BMP4. BMP signaling activity is restricted to the edge of the colony over time where it causes differentiation to cells that most closely resemble the amnion^17^. Simultaneously, BMP activates transcription of Wnt ligands, and Wnt signaling in turn activates transcription of Nodal. This causes Wnt and Nodal signaling activities to spread inward towards the colony center with the front of Wnt signaling starting earlier, but the front of Nodal signaling moving faster. These fronts provide the cues for mesendoderm differentiation.

Although much has been learned from this model, the presentation of a high dose of BMP is an artificial feature that hinders understanding the communication between amnion and epiblast. To overcome this limitation, here we created a model in which cells are pre-differentiated to amnion-like cells (AMLCs) and then juxtaposed with epiblast cells without adding differentiation signals to the media. We show that this juxtaposition is sufficient to trigger the formation of a primitive-streak like region in the epiblast cells that border the AMLCs. Cells express markers of primitive streak and then undergo directed migration away from the AMLC-epiblast border. These primitive streak cells then differentiate into endoderm and several mesodermal subtypes including axial, paraxial, and lateral mesoderm. The BMP-Wnt-Nodal signaling hierarchy is preserved in this model with signaling fronts that begin at the amnion-epiblast border and move into the epiblast territory. Performing knockouts in both the epiblast and amnion-like compartments revealed more intricate feedback between these tissues than previously appreciated. In particular, Nodal ligand in both the epiblast and amnion plays important roles in establishing appropriate patterning in the epiblast. Taken together, our study establishes a system to study epiblast-amnion communication and shows that feedback between these tissues is important for controlling signaling dynamics and patterning at gastrulation.

## Results

### Juxtaposition with AMLCs is sufficient to induce mesendoderm differentiation in hPSCs

hPSCs treated with BMP ligands together with Activin/Nodal inhibitors differentiate to nearly pure populations of extraembryonic cells with expression of CDX2, ISL1, GATA3, and Hand1^2,16–21^. We and others have previously shown that these cells are most similar to amnion cells in the primate embryo, consistent with their origin from hPSCs, which are most similar to epiblast^16,17^. Moreover, studies with naïve hPSCs have shown that differentiation to amnion, but not trophectoderm, is dependent on BMP signaling^20^. We refer to these cells as amnion-like cells (AMLCs).

We used AMLCs to create a model for the interactions between amnion and epiblast that occur at the embryonic-extraembryonic border (Figure 1A). We pre-differentiated cells labeled with a fluorescent H2B fusion protein to AMLCs. We initially simply mixed these AMLCs with hPSCs and found that the two populations segregated in time and that there was sporadic expression of BRACHYURY (BRA) within the hPSC population (Figure S1A), indicating that AMLCs may be capable of inducing differentiation in hPSCs.

**Figure 1.**
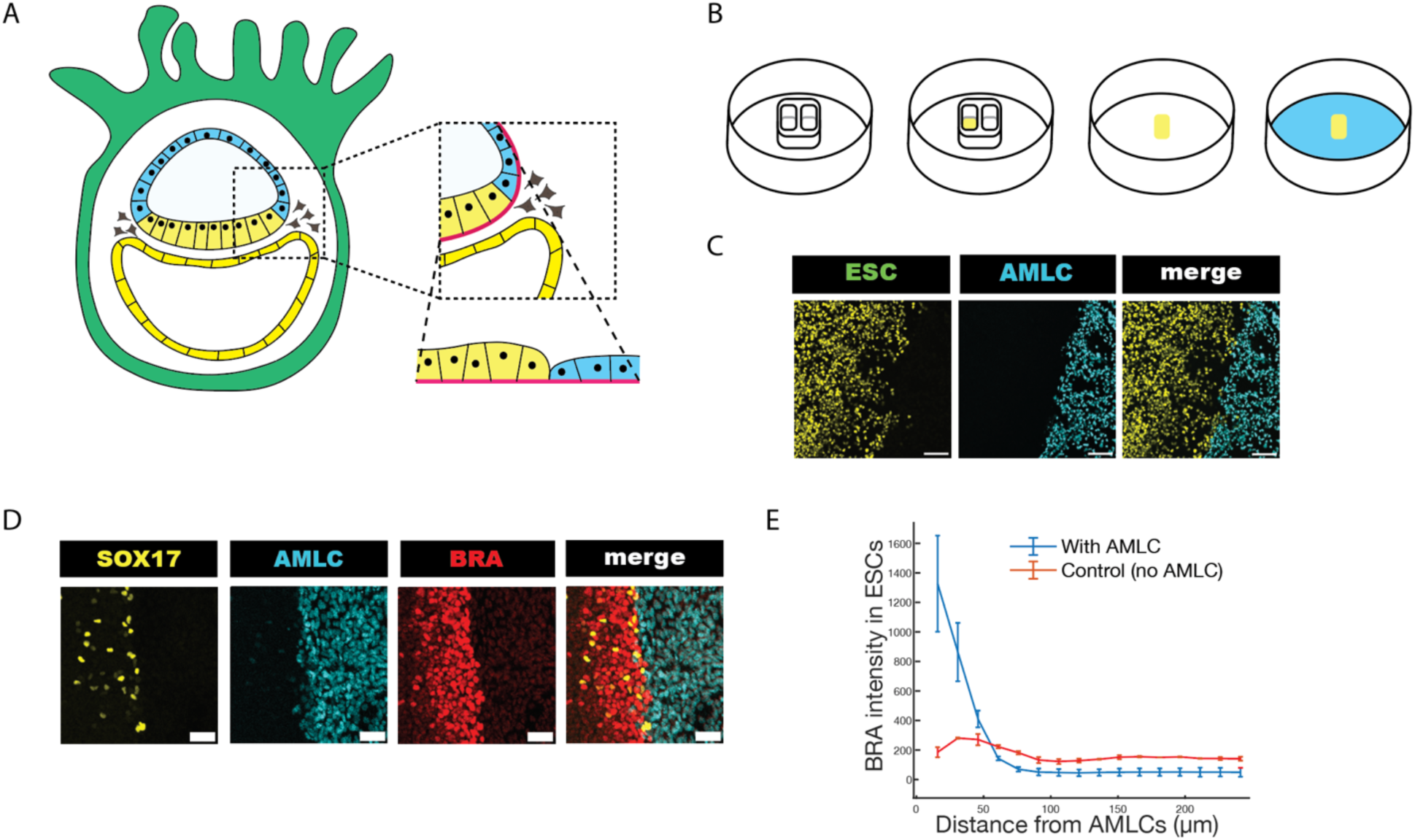
Amnion-like cells (AMLCs) induce mesoderm and endoderm differentiation in human embryonic stem cells (ESCs). (A) Schematic of the human embryo at the onset of gastrulation indicating the region modeled in the study. (B) Schematic of the experimental procedure. (C) Representative images showing the border created between hPSCs and AMLCs. hPSCs are labeled by Venus::H2B while AMLCs are labeled by Cerulean::H2B. (D) Representative images of immunofluorescence for markers of mesoderm (BRA; red) and endoderm (SOX17; yellow) differentiation at the AMLC-hPSC border after 60h of interaction. (E) Quantification of BRA intensity as a function of distance from the border between cell types. Error bars represent standard error of the mean (s.e.m). Quantifications represent averages over 5 independent images. Scale bars: 100 μm (C), 50 μm (D).

To study this hPSC-AMLC interaction in a more controlled and quantitative manner, we developed a protocol using a removable cell culture insert (Figure 1B). We used the insert to seed hPSCs in a defined area, removed the insert, and then seeded AMLCs which had been differentiated from hPSCs for two days. The AMLCs adhere to surrounding area but not on top of the hPSC population, creating a defined border between the two populations (Figure 1C). After 60 hours of juxtaposition, the hPSCs adjacent to the AMLC domain express BRA, a marker of primitive streak and mesoderm differentiation, and SOX17, a marker of endoderm (Figure 1D). The robustness of mesendoderm differentiation is heavily dependent on the seeding density of the AMLCs (Figure S1C). When AMLCs are seeded at low densities (20,000 cells/cm^2^), the border between the hPSC colony and the AMLCs is not well defined, and there is no differentiation in the hPSCs. At medium seeding densities of 40-60,000 cells/cm^2^, the hPSC colony adopts a more defined shape and patches of differentiation marked by BRA are found near the boundary. At high seeding densities of 150,000-200,000 cells/cm^2^, there is a continuous ribbon of BRA positive cells at the border. In contrast, hPSCs do not differentiate when seeded using an identical protocol but without seeding AMLCs surrounding them or when AMLCs are substituted with previously described human trophoblast stem cells^22^ (Figure 1E, Figure S1B).

### Juxtaposition with AMLCs induces gastrulation-like cell migration and patterning

To understand the dynamics of AMLC-induced hPSC differentiation, we observed BRA and SOX17 expression after 1-4 days of juxtaposition (Figure 2A,B) with immunostaining. BRA expression begins with a band of differentiation a few cell diameters wide on day 2 at which time there are scattered cells within this band expressing SOX17. On day 3, BRA intensity increases, and the band of differentiation becomes wider, with most BRA+ cells found in this band and a minority found migrating away from it. SOX17 expression can also be found both inside the band and in isolated cells farther from the border than the BRA-positive cells. On day 4, the expression of BRA and SOX17 both expand dramatically with greater numbers of cells appearing outside the primitive streak-like band near the hPSC-AMLC boundary. This pattern is consistent with the patterns of BRA and SOX17 observed in vivo with expression of BRA in the primitive streak preceding that of SOX17, with the primitive streak expanding away from the amnion as gastrulation proceeds, and with SOX17 expressing cells eventually migrating further anteriorly than BRA expressing cells.

**Figure 2.**
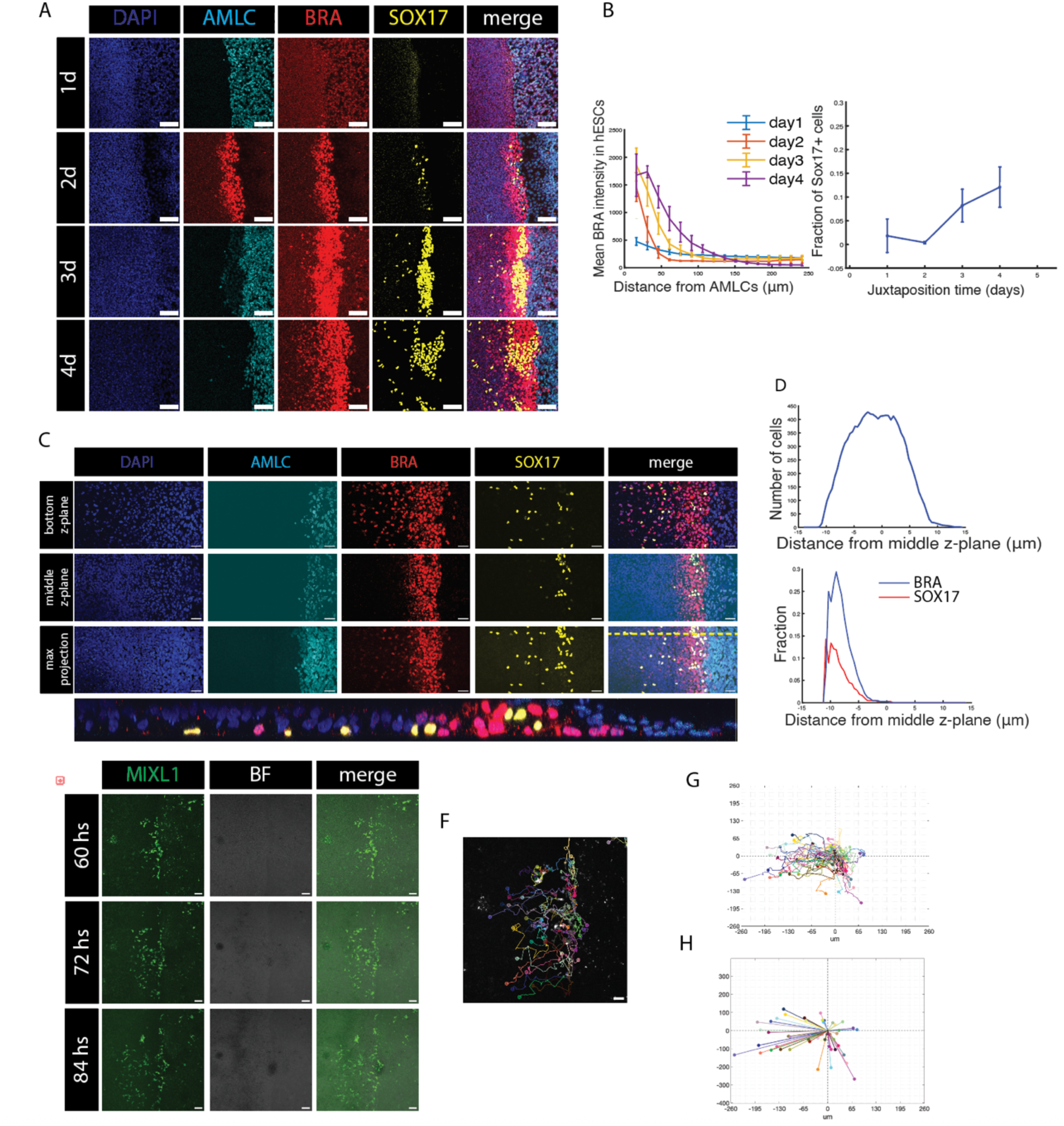
Differentiated cells undergo directed migration away from the border. (A) Representative immunostaining of BRA and SOX17 after 1-4 days of juxtaposition. (B) Quantification of average BRA intensity and fraction of SOX17 positive cells as a function of the distance to the AMLCs. (C) Images of immunostaining at different z-planes (plane interval 0.46 um) after 3.5 days of interaction. (Bottom) Orthogonal view taken at the yellow dash line (bottom-right) with the plate surface at the bottom. (D) Quantification of the total number of cells detected in each z-plane (top) and the fraction of BRA/SOX17 positive cells (bottom), plotted as a function of the distance to the middle plane. (E) Snapshots of a live imaging experiment using MIXL1 reporter cells diluted 1:20 in unlabeled hESCs at the indicated times after the cell types were juxtaposed. (F) An image from the movie in D taken at 60 h overlaid with tracking trajectories of 46 MIXL1 positive cells from 60 h (as asterisks) to 84 h (as circles). Scale bar, 50 um (A and C), 100 um (E and F). (G) Single cell migration trajectories shown in E and F with each starting position overlayed at (0,0). (H) Single cell displacement shown in E and F with each starting position overlayed at (0,0).

We investigated the three-dimensional structure of the boundary region at day 3.5 of juxtaposition (Figure 2C). At this point, the primitive-streak like region becomes three dimensional with BRA+ cells enveloping SOX17+ cells. Away from this region, the hPSCs retain their epithelial character while the BRA+ and SOX17+ cells are found beneath this layer, consistent with the positions of migrating mesendodermal cells in vivo.

To determine whether the mesendodermal cells induces by AMLCs differentiate further, we examined a panel of markers after 4 days of juxtaposition (Figure S2), as this is the latest point before more extensive spatial spreading of mesoderm cells may complicate interpretation of the resulting patterns. In addition to BRA and SOX17, we detected MIXL1, an additional primitive streak marker, GATA6, an additional endoderm marker, as well as markers of paraxial (TBX6), axial (GOOSECOID, FOXA2), and lateral mesoderm (HAND1, ISL1) (Figure S2, S3). We also observed occasional HAND1+ cells that were derived from the epiblast layer but migrated underneath the amnion, likely consistent with extraembryonic mesoderm identity. Thus, the primitive streak cells resulting from the juxtaposition of amnion and epiblast give rise to endoderm as well as all the major subtypes of mesoderm.

To understand whether the differentiated cells found at a distance from the border result from cell migration from the border region, we performed live imaging with an established MIXL1 reporter cell line^23^. Live imaging showed MIXL1 intensity increasing on day 2 with the cells expressing it remaining largely stationary near the border with the AMLCs. On day 3, a portion of the MIXL1 expressing cells move away from the border. To track individual migrating cells, we diluted MIXL1 reporter cells 1:20 in unlabeled human H9 cells, used this mixture in the hPSC compartment in the juxtaposition assay, and performed live imaging. To capture the migration, we chose the window from 60 to 84 hrs (Figure 2E). We used the tracking module in ilastik^24^ to track 46 individual MIXL1 expressing cells through this time interval. To visualize the directionality of the cell migration, we aggregate all 46 migration tracks to a single starting point (Figure 2G,H), the tracks show a clear bias branching away from the AMLC-hPSC border into the epiblast region, with a smaller number of cells migrating into the amnion region, potentially consistent with migration patterns of extraembryonic mesoderm (Figure 2E-H).

### Endogenous BMP, Wnt, and Nodal signaling are required for juxtaposition-induced patterning

To assess the involvement of endogenous BMP, Wnt and Nodal signaling, we inhibited these signaling pathways with Noggin, IWP2, and SB431542 (SB), respectively. Loss of signaling through any of these pathways caused a dramatic decrease in primitive streak and endoderm differentiation as reflected in the markers BRA, MIXL1 and SOX17 (Figure 3). This suggests that, as in the mouse and in micropatterned hPSC colonies, signaling activity through all three of these pathways is required for gastrulation initiation. Interestingly, treatment with each inhibitor also altered the morphology of AMLCs, suggesting ongoing signaling regulated the behavior of AMLCs as well as the hPSCs.

**Figure 3.**
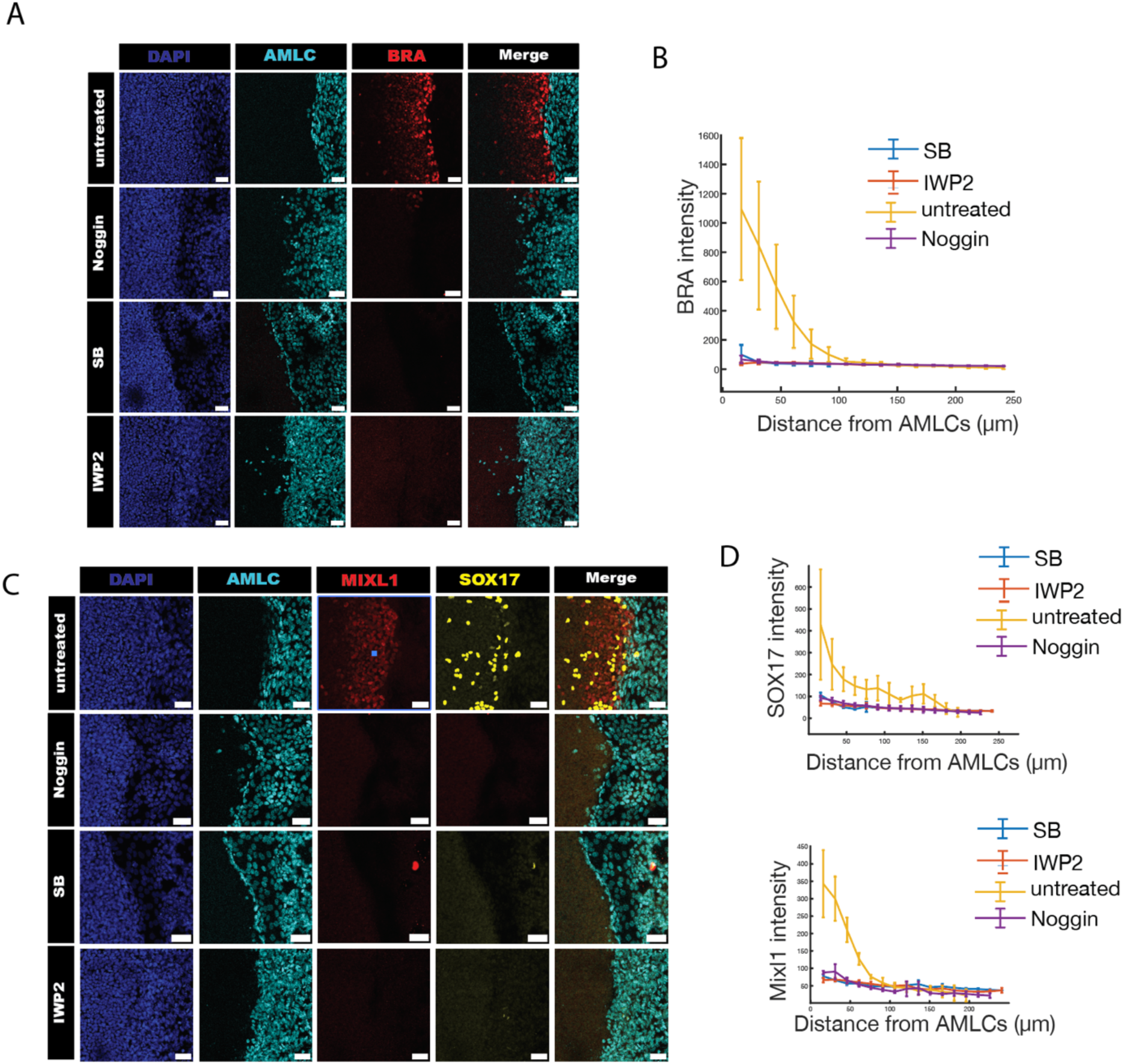
BMP, Wnt and Nodal signaling are required for the patterning induced by AMLCs. (A) Representative immunostaining of BRA after 3.5 days of interaction with BMP, Nodal, or Wnt signaling inhibited by Noggin, SB and IWP2 respectively. (B) Quantification of average BRA intensity as a function of the distance to AMLCs. (C) Representative immunostaining of MIXL1 and SOX17 with BMP, Nodal or Wnt signaling inhibited. (D) Quantification of average SOX17 and MIXL1 intensity as a function of the distance to AMLCs. Error bar represents s.e.m. Scale bars, 50 μm.

To understand the signaling dynamics that underlie patterning, we performed a time course from day 1 to day 3 of juxtaposition with immunostaining for pSMAD1, SMAD2 and BRA (Figure 4A-D). BMP signaling activity, as reflected in pSMAD1 intensity, shows a clear gradient within the epiblast compartment by day 1, before any expression of differentiation markers. The pSmad1 gradient expanded in both intensity and spatial extent between days 2 and 3, during which time BRA expression was initiated close to the boundary and expanded. In contrast,

**Figure 4.**
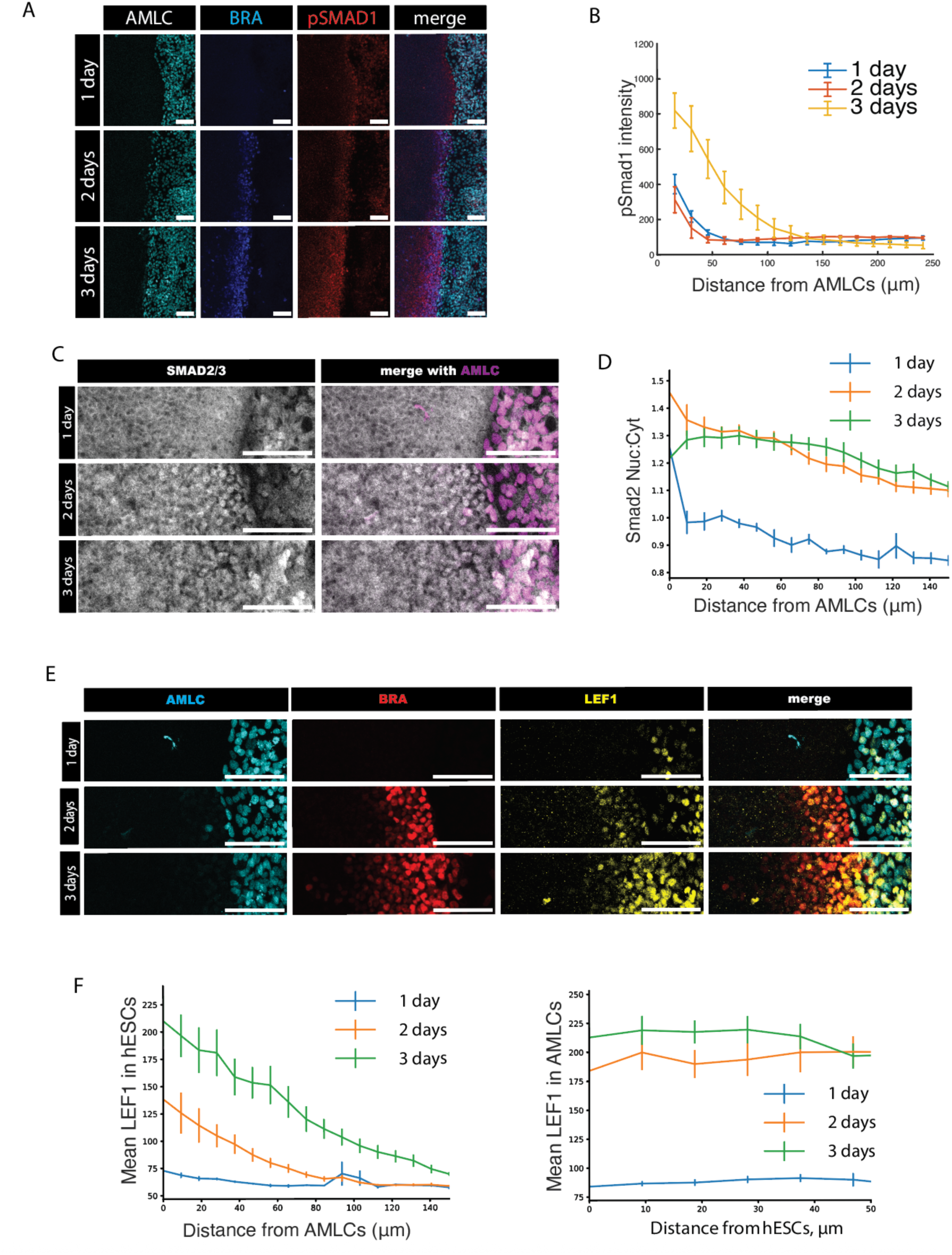
Dynamics of BMP, Nodal, and Wnt signaling activity in the juxtaposition experiment. (A, C, E) Representative immunostaining of pSMAD1 (A), SMAD2 (C), and LEF1 (E) after 1, 2, or 3 days juxtaposition as indicated. In A and E, BRA is also co-staining. Scale bar, 100 μm. (B, D, F) Quantification of pSMAD1 intensity (B), SMAD2 nuclear to cytoplasmic ratio (D), and LEF1 (F) in the hESCs after 1, 2, or 3 days post-juxtaposition plotted as a function to their distance to the AMLCs. In F, Quantification of LEF1 in the amnion is also shown (right panel). Error bars are standard error of means (s.e.m.), n > 5 images for all experiments.

Nodal activity as reflected in SMAD2/3 nuclear localization was not present in the hPSC compartment until day 2 and spread into the hPSC region, following the same trend as BRA. Within the AMLC compartment, BMP activity is the strongest on day 1 and progressively decreases through day 3 and 4, while Nodal activity remains strong through day 4 (Figure S4). The timing of BMP activity suggest that it is the primary signaling cue that initiates gastrulation-like events in this model of human amnion-epiblast communication.

To investigate the dynamics of Wnt signaling, we used immunostaining for the direct Wnt target LEF1 (Figure 4E,F). WNT activity began near the boundary with the AMLCs and was slightly elevated by 24 hours. Over the next 24 hours, signaling near the boundary intensified while the activity spread inward reaching approximately 100μm from the boundary. The timing of Wnt signaling is consistent with Wnt acting as an intermediary between BMP and Nodal signaling as has been observed in other systems. Within the amnion compartment, Wnt signaling rose dramatically between day 1 and day 2 concomitant with its spread through the epiblast compartment and remained at high levels through day 3 (Figure 4F).

### BMP is upstream, while Nodal is downstream, of Wnt signaling

To better understand the hierarchy of signaling that patterns the hPSC at the border with AMLCs, we inhibited each pathway with Noggin, SB or IWP2 (Figure 5A,B) and stained for SMAD2/3. As expected, SB effectively inhibited Nodal throughout the hPSCs and minimal SMAD2/3 was found in the nucleus. Both Noggin and IWP2 treatment dramatically lowered nuclear Smad2/3 in the hPSCs, placing Nodal signaling downstream of both BMP and Wnt in this model. Interestingly, IWP2 did not affect SMAD2/3 localization in AMLCs, suggesting Nodal signaling in this compartment is independent of Wnt (Figure S4B).

**Figure 5.**
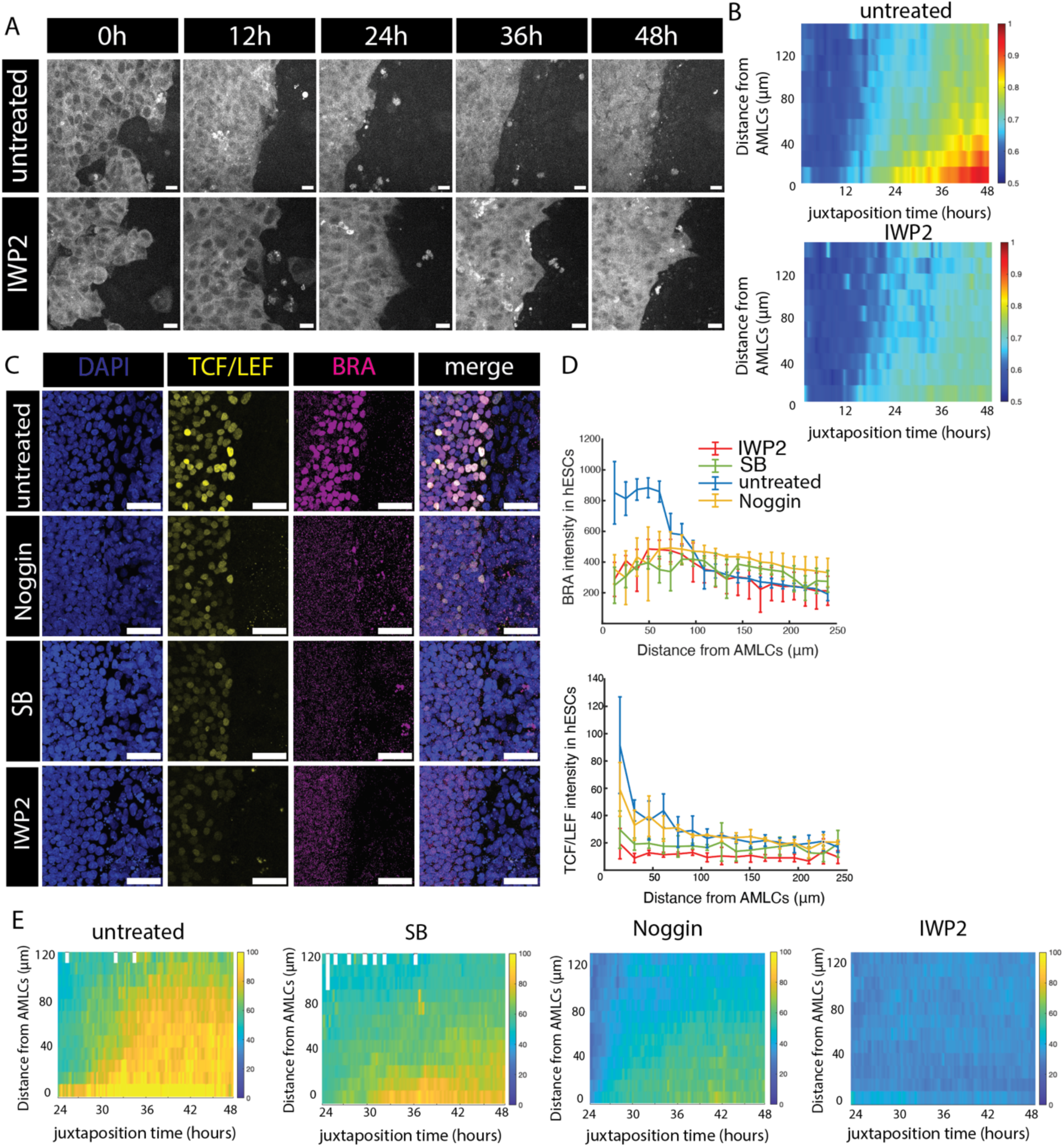
Signaling dynamics in the juxtaposition model. (A) Snapshots taken at the indicated times from a live imaging experiment using GFP::SMAD4 reporter cell line. (B) Quantification of SMAD4 nuclear:cytoplasmic ratio in the live imaging experiment shown in A, plotted as a color-coded kymograph with time as the x-axis and distance from the closest CFP-labeled AMLC cell as the y-axis. (C) Immunostaining of BRA (magenta) and TCF/Lef::GFP (green) after 2 days of juxtaposition, with BMP, Nodal, or Wnt signaling inhibited by Noggin, SB and IWP2, as indicated. (D) Quantification of Wnt activity (TCF/Lef intensity) and BRA expression as functions of the distance from the nearest AMLCs. Error bar, standard error of means (s.e.m.), n > 5. (E) Quantification of TCF/LEF intensity from live imaging plotted as a color-coded kymograph with time as the x-axis and distance from the closest CFP-labeled AMLC cell as the y-axis.

We used a SMAD4 reporter cell line^12,25^ to report on Nodal and BMP signaling during the first 48 hours of the interaction (Figure 5C,D). SMAD4 starts to localize to nuclei within the hPSC compartment between 12 and 24 hours after juxtaposition and this early signaling was not blocked by addition of IWP2 to inhibit WNT signaling. Together with the pSmad1 data above, this data suggests that BMP signaling in the epiblast compartment begins during the first day of juxtaposition and does not require WNT signaling, consistent with BMP acting as the most upstream signal. After 24 hours, nuclear SMAD4 spread inward from the border and this increase was mostly blocked by IWP2. Together with the SMAD2 data above, this suggests that the spread of Nodal signaling which occurs on day 2 is downstream of Wnt signaling and is blocked by Wnt inhibition.

To further examine the position of Wnt signaling within this hierarchy, we examined the dynamics of the TCF:LEF reporter with each of the inhibitors (Figure 5E, S6). As expected, IWP2 treatment, which prevents the secretion of WNT ligands, completely blocked upregulation of the reporter. Inhibiting BMP signaling also blocked nearly all signaling, consistent with BMP acting upstream of Wnt. In contrast, inhibiting Nodal with SB only slightly reduced WNT activity as reflected in the TCF:LEF reporter.

We also measured Wnt activity using LEF1 immunostaining (Figure S5). Consistent with the reporter data Noggin and IWP2 both strongly reduced LEF1 expression. SB strongly increased LEF1 expression, in contrast to the minor reduction observed in the TCF:LEF reporter. This could either reflect complexity in the regulation of different Wnt targets or a separate inhibitory effect of Nodal on LEF1 protein expression. Within the amnion, we also observed strong inhibition of LEF1 with IWP2 but opposite trends with SB and Noggin – SB decreased and Noggin increased LEF1 expression. Taken together, our data are consistent with the BMP, Wnt, Nodal hierarchy, which has previously been described in the mouse and in micropatterned colonies, operating in the juxtaposition model as well, but they also reflect additional complexity in that this hierarchy does not operate within the amnion compartment.

### Human gastrulation requires robust reciprocal Nodal signaling between amnion and epiblast

An advantage of the juxtaposition model is the ability to using genetically modified cells in only the epiblast or only the amnion compartment, analogous to tissue specific knockouts in vivo. We used previously described NODAL-KO cells^2^ in either the epiblast, the amnion or both compartments. Consistent with an essential role for epiblast derived Nodal in gastrulation, we found loss of Nodal in the hPSCs caused a near complete loss of mesendoderm differentiation regardless of whether NODAL-KO cells were used in the AMLC compartment. Interestingly, Nodal from the amnion was required to achieve full differentiation as using NODAL-KO cells in the amnion compartment caused a reduction in BRA expression and a loss of endoderm marker SOX17 (Figure 6A,B).

**Figure 6.**
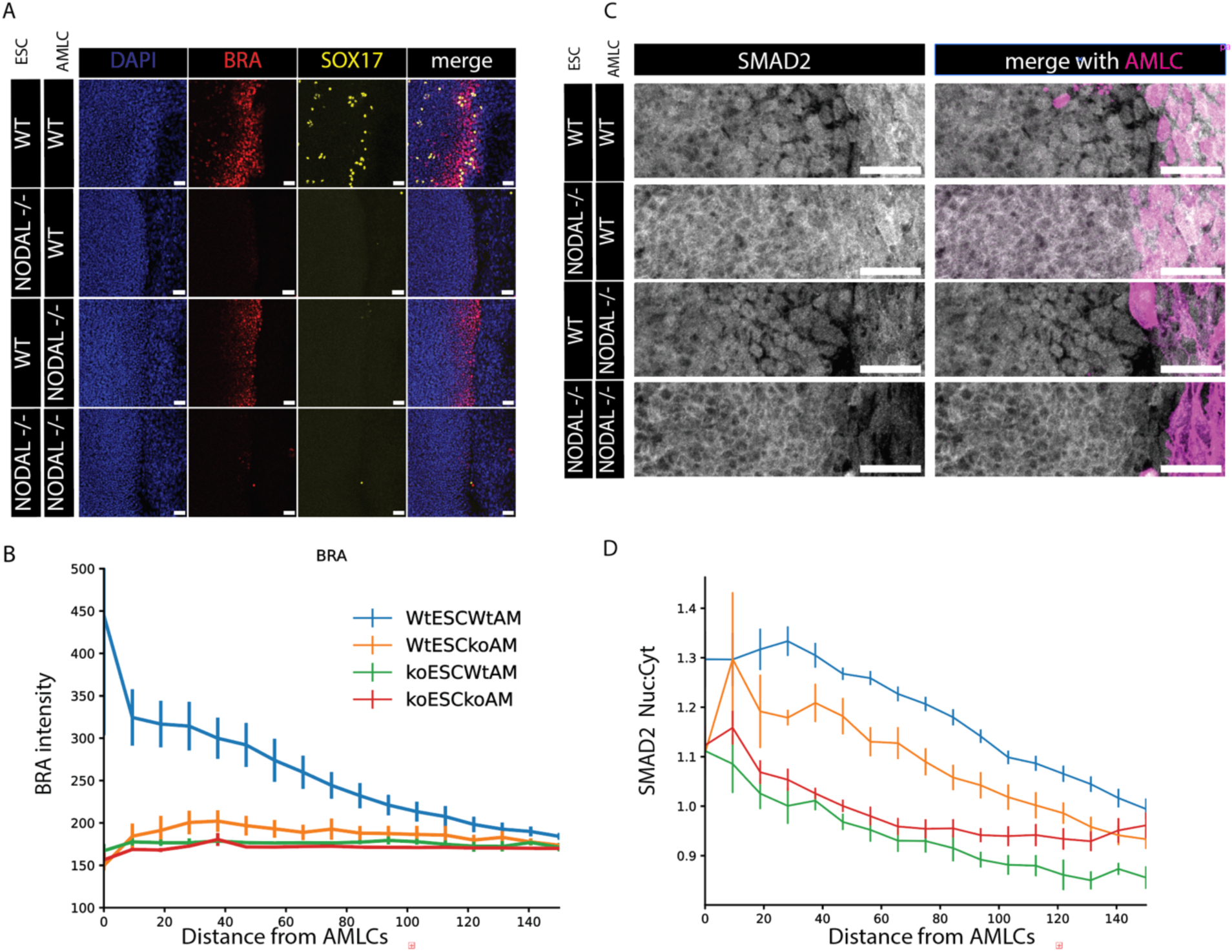
Reciprocal Nodal signaling between epiblast and amnion is required for endoderm differentiation in the juxtaposition model. (A, B) Representative images (A) and quantification (B) of fate markers BRA and SOX17 for juxtaposition experiments performed with cells with the indicated genotypes. (C, D) Representative images of SMAD2 (C) and quantification of the SMAD2 nuclear:cytoplasmic ratio (B) for juxtaposition experiments performed with cells with the indicated genotypes. Scale bars 100um, n>5 images for all conditions.

To better understand the role of Nodal in each compartment, we stained each of these combinations for SMAD2/3 (Figure 6C,D). Consistent with the above, loss of NODAL in the epiblast compartment strongly compromised signaling within this compartment, regardless of which cells were used in the AMLC comportment. Loss of NODAL in the amnion compartment led to a more mild reduction of signaling in the epiblast, consistent with the diminished, but still present, mesoderm differentiation.

### Amnion is a potential precursor for extra-embryonic mesenchyme

Finally, we asked whether the ability of amnion to induce gastrulation behaviors is a stable or transient property. We pre-differentiated cells to amnion fates for varying lengths of time and then used these in the juxtaposition assay. We found that cells differentiated for 2 or 3 days are capable of inducing BRA expression in the epiblast compartment, while cells treated for longer or shorter are not (Figure 7A,B). Examining markers of amnion fates, we found that the loss of the ability to induce BRA in the epiblast correlated with a reduction in amnion markers such as CDX2 and ISL1 (Figure 7C,D). We also found that as cells began to downregulate CDX2 and ISL1 they upregulated VIM, a key marker of extraembryonic mesenchyme (Figure 7C,D).

**Figure 7.**
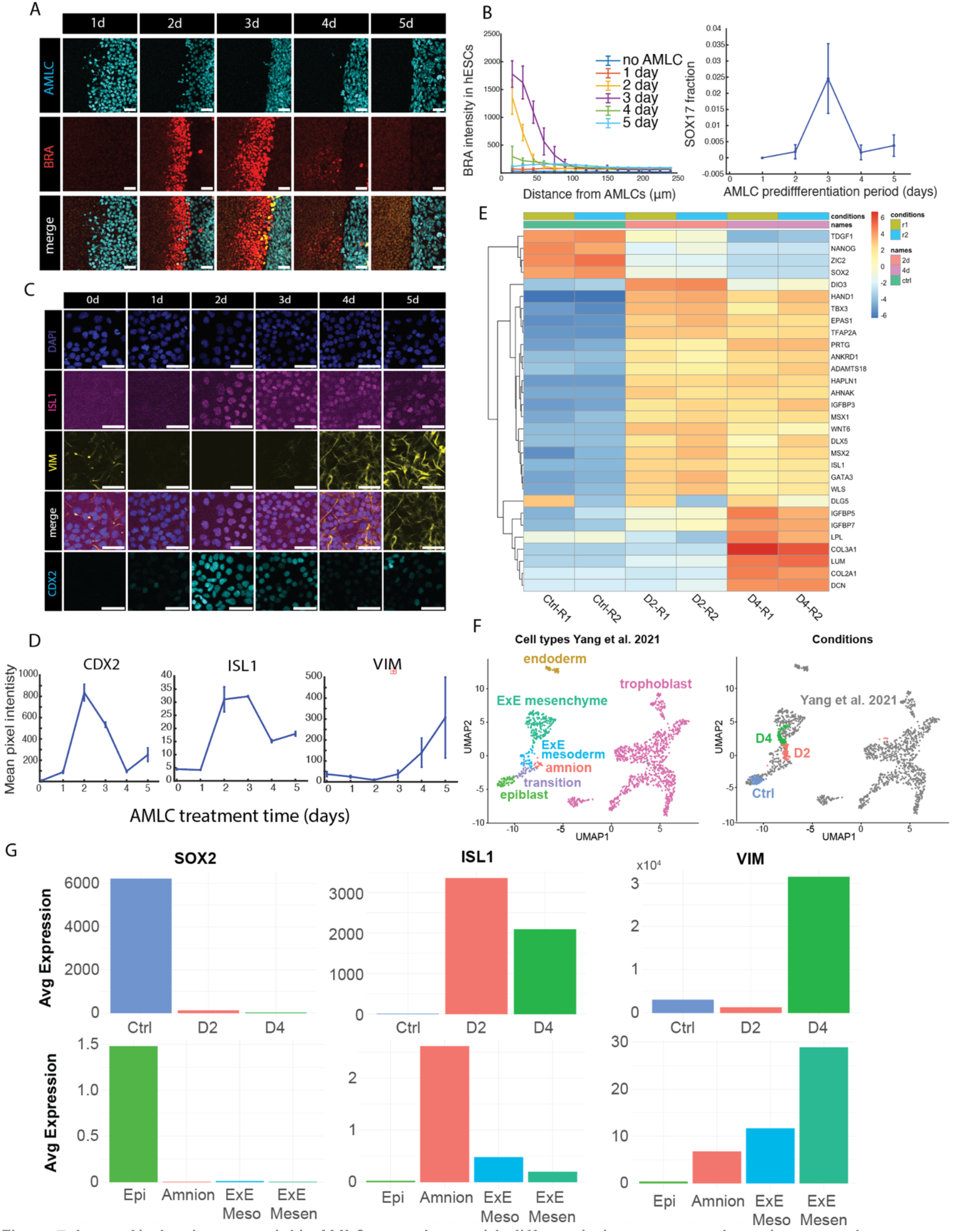
Loss of inductive potential in AMLCs correlates with differentiation to extraembryonic mesenchyme. (A) Immunostaining of BRA from a juxtaposition experiment where the timing of pre-differentiation was varied from one to 5 days. (B) Quantification of BRA and SOX17 from the experiment in (A). (C) Immunostaining of pluripotent cells grown in AMLC differentiation media for 0 to 5 days. (D) Quantification of ISL1, CDX2 and VIM from the experiment in (C) (E) Quantification of pluripotent, amnion, and extraembryonic mesenchyme markers in RNA sequencing data from cell differentiated for 0, 2 or 4 days. (F) (left) Single cell RNA sequencing data from non-human primates with cell types indicated. Data from Yang et al.^10^ (right) Data from Yang et al (gray dots) combined with data from differentiation of hPSCs to amnion (control, d2 and d4). (H) Average expression of key markers during amnion differentiation in vitro (top row) and in cell types in the non-human primate embryo (bottom row).

To better understand the differentiation of the AMLCs, we performed RNA sequencing after 2 or 4 days of differentiation. These data showed that in addition to CDX2 and ISL1, other amnion markers including GATA3 and TFAP2A/C peak on day 2 and decline by day 4 while markers of extraembryonic mesenchyme including GATA4, GATA6, LUM, COL2A1, and COL3A1 are upregulated on day 4.

To analyze these data more globally, we integrated them with existing scRNA sequencing for cynomolgus monkey (Yang et al^10^, Figure 7F, G) and a human complete embryo model (Oldak et al^26^, Figure S7). In UMAP projections of the data from Yang et al, amnion cells can be found adjacent to epiblast cells with cells classified as extraembryonic mesenchyme or extraembryonic mesoderm also found within the same large group of cells and cells of the trophectoderm found in a well-separated cluster (Figure 7F). We used a method for down-sampling our bulk RNA sequencing data to create pseudo single-cell data^27^ and mapped these to the same space as the monkey embryo data. The untreated control cells mapped to the epiblast, the day 2 differentiated cells to the amnion, and the day 4 cells to a region containing both extraembryonic mesoderm and mesenchyme (Figure 7F). The expression of key marker genes was consistent between the day 2 differentiated cells and the amnion and the day 4 differentiated cells and the extraembryonic mesenchyme. The mapping to the human embryo model data produced similar results with control, day 2, and day 4 cells mapping to regions annotated as epiblast, amnion, and “ExEM-like” cells, respectively (Figure S7).

## Discussion

In this study, we modeled the initiation of gastrulation at the embryonic-extraembryonic boundary at the onset of gastrulation by juxtaposing hPSCs and AMLCs. We show that this juxtaposition is sufficient to trigger events in the hPSCs which are associated with gastrulation in vivo including differentiation to primitive steak, subsequent diversification of mesendodermal fates, and directed cell migration away from the boundary, which corresponds to anterior-ward migration in this culture system. The hierarchy of BMP, Wnt, and Nodal signaling previously observed in mammalian model organisms in vivo^1^ and human gastrulation models in vitro^2,28^ is preserved in the hPSC compartment, however, this cascade does not operate in the AMLC compartment. Further, while BMP signaling originating from the amnion is the trigger for the gastrulation-like events, generating the full spectrum of cell fates requires ongoing reciprocal signaling between the two compartments. In particular, we show that the Nodal ligand is required in both compartments to generate a pattern containing both mesoderm and endoderm. Finally, we show that the ability of AMLCs to induce the events of gastrulation in the hPSCs is a transient property that is acquired as amnion cells differentiate from the epiblast. The loss of inductive ability correlates with the gain of mesenchymal properties. Transcriptomic analysis shows that these cells correspond to extraembryonic mesenchyme seen in complete human embryo models as well as primate embryos, suggesting the amnion may be one source of these cells in vivo.

Classic embryological experiments in the mouse have shown that the process of gastrulation relies on intimate coordination between the epiblast and the extraembryonic tissues both to initiate primitive streak formation and to restrict it to the posterior side of the embryo. BMP4 from the trophectoderm is essential to this process^3^, however, the geometry of primate and rodent embryos differ substantially^7,9,29^. In the mouse, at the onset of gastrulation, the trophectoderm and epiblast are juxtaposed and surround a single cavity, while the amnion does not form until later. In contrast, in primates, the amnion forms before gastrulation and it is the amnion that surrounds a single cavity with the epiblast, which has led to the idea that interactions between the amnion and epiblast may be the trigger for gastrulation. Our data here show directly that juxtaposition of amnion-like cells with hPSCs in vitro is sufficient to trigger many of the hallmarks of embryonic gastrulation.

BMP signaling originating from the extraembryonic tissues initiates mesendoderm differentiation by transcriptionally activating the Wnt3 ligand in both the visceral endoderm and the epiblast^30,31^. Wnt signaling in turn transcriptionally activates the Nodal ligand, and all three of these ligands are essential for gastrulation. These connections were originally established in the mouse and have also been validated in micropatterned hPSC colonies treated with BMP4. We show here that the same cascade operates in the hPSC compartment of the juxtaposition model. Measurements of signaling dynamics from micropatterned hPSCs show that both Wnt and Nodal spread through the colony covering an increasing area of the colony in time. Our measurements here establish similar dynamics in the juxtaposition model, showing that these dynamics are not an artifact of the exogenous BMP stimulation, and that a tissue boundary alone is sufficient to trigger these dynamics, which can then serve to pattern the variety of cell fates that emerges during gastrulation.

In mouse, Nodal ligand from the epiblast is important for establishing and maintaining the source of BMP in the trophectoderm, however, a requirement for Nodal expression in this tissue has not been documented. In contrast, we show here that Nodal is required in both the amnion and epiblast compartments to achieve full patterning in the juxtaposition model. Loss of Nodal in the amnion leads to a loss of endoderm differentiation in the epiblast. Nodal signaling in the epiblast is also reduced by loss of the ligand in the amnion, which likely explains the loss of endoderm differentiation. It is however unlikely that Nodal ligand from the amnion directly signals within the epiblast, as we previously showed that Nodal only functions at very short range in this context. Instead, reciprocal signaling at the extraembryonic-embryonic boundary is needed to strongly activate a transcriptional relay involving Nodal activation of its own expression. Consistent with this interpretation, juxtaposition of Nodal-KO cells with Nodal expressing cells also reduces Nodal signaling in the expressing cells despite the limited range of the ligand^32^.

Gastrulation in all amniotes is associated with an EMT in the primitive streak and migration of the resulting cells in the anterior direction. In the juxtaposition model, cells transit through the primitive streak like region beginning at the tissue boundary and migrate into the epiblast compartment, underneath the original layer of cells, and away from the boundary. Cells show directed migration that persists for at least 100-200um from boundary. How cells maintain this sense of direction over long time durations and distances is an interesting question for future study.

In primate embryos, extraembryonic mesenchyme (ExMC) emerges before gastrulation and forms a layer in between the trophectoderm and the remainder of the embryo. Several sources have been suggested for these cells including the visceral endoderm and the trophectoderm^33,34^. Here we show that as the cells of the amnion lose the ability to induce differentiation in the epiblast, they can further differentiate to mesenchyme which resembles the ExMC. These possible sources of ExMC are not mutually exclusive, and it is likely that these cells derive from multiple sources in order to surround the embryo.

The model developed here has several advantages over previously developed gastrulation models. Once the AMLCs are differentiated, it requires no addition of exogenous ligand, allowing the observation of reciprocal signaling in a more natural context. Moreover, the use of separate cell populations in the amnion and epiblast compartments makes it straightforward to use knockout cells in only one compartment or the other, which facilitates the dissection of signaling interactions between compartments. This model should serve to help understand how interactions between the human embryo proper and the surrounding tissue initiate gastrulation and drive pattern formation during this critical developmental stage.

## Materials and Methods

### Cell Lines

All experiments were conducted using the hESC line ESI017 unless otherwise specified. The MEL1-MIXL1 reporter cell line^23^, used in tracking experiments, and the TCF/LEF reporter cell line^35^ were generous gifts from Ed Stanley (MCRI, Australia) and Jianping Fu (University of Michigan), respectively. SMAD signaling dynamics were monitored using a previously published GFP::SMAD4 reporter derived from RUES2 cells^25^. Nodal and Lefty knockout lines were generated from ESI017 cells and characterized in previous studies from the laboratory^2,32^.

### Routine Cell Culture

Human PSCs were maintained in mTeSR1 medium (StemCell Technologies) under standard conditions (37 °C, 5% CO₂). Cells were cultured on dishes coated with Matrigel or Geltrex (dilution 1:400 in DMEM/F12). Routine passaging was performed using dispase (StemCell Technologies), and cultures were regularly tested and confirmed negative for mycoplasma contamination.

### AMLC differentiation

**A** single cell suspension of hPSCs was collected using accutase and seeded at a density of 40,000 cells/cm^2^ in a Matrigel coated dish containing mTeSR1 supplemented with 10 µM ROCK inhibitor Y-27632, 50 ng/ml BMP4 and 10 µM SB431542. The seeded dishes were then incubated at 37 °C with the treatment refreshed every 24 hours.

### Juxtaposition Assay

A single cell suspension of hPSCs was collected using accutase. The concentration of hPSCs was adjusted to 2.5-3 x 10^6^ cells/ml. to keep the seeding volume at the optimal 70-75 µl. Then a single removable insert (ibidi #80209) was planted at the center of a Matrigel coated 8 well slide (ibidi #80806). After confirming the insert was firmly attached to the slide, 150,000 hPSCs were seeded into the insert chamber. This slide was then incubated at 37 °C for 1 hour. Following incubation, the insert and media were removed, and the seeded cells were gently washed with PBS. Cells were then incubated overnight in mTeSR1 medium. On the following day, pre-differentiated AMLCs were seeded onto the hESCs and incubated in mTeSR1 medium supplemented with 10 μM ROCK inhibitor Y-27632 (RI) for 2 hours. After this incubation, the medium containing RI was removed, fresh mTeSR1 was added, and the culture was maintained in mTeSR1 for the remainder of the experiment. The medium was refreshed every 24 hours.

Each experiment was accompanied by a control group that did not contain AMLCs. In this control group, to mimic the physical border created by juxtaposition in the experimental group, the plate was coated differently. Specifically, the insert was placed on an uncoated surface, and 70 μL of Matrigel mix was added inside the insert well. This configuration ensured that hPSCs could not grow beyond the coated region, establishing a defined boundary.

### Immunofluorescence Staining

Cells were fixed with 4% paraformaldehyde for 20 minutes at room temperature. Following fixation, samples were permeabilized and blocked for 30 minutes in a blocking solution containing 3% donkey serum and 0.1% Triton X-100 in PBS-/-. Primary antibodies were diluted in blocking solution and incubated with samples overnight at 4 °C. The next day, samples were washed three times in PBST (0.1% Tween-20 in PBS), each for 20 minutes at room temperature. Secondary antibodies, with 2.5 ng/mL DAPI included, were added in blocking buffer and incubated for 30 minutes at room temperature in the dark. After removal of the secondary antibody solution, samples were washed twice in PBST and stored in PBS at 4 °C until imaging. A list of antibodies and dilutions is included in Table S1.

### Imaging

Fixed samples were imaged using an Olympus IX83 laser scanning confocal microscope with a 10×, NA 0.4 objective unless otherwise indicated. Higher magnifications were used for specific figures: Figure 2C (40× Si oil objective) and Figure 7C (20× NA 0.75 objective). Live imaging of GFP::SMAD4 reporter cells (Figure 4G) was performed on an Olympus/Andor spinning disk confocal microscope using a 40× Si oil objective.

### Image processing

Images of the border between AMLCs and hPSCs were used to create a mask of the AMLCs with ilastik. A distance transform was used to determine the distance of each pixel to the border and then averages were determined as a function of this distance. For experiments involving SMAD2, it was necessary to perform calculations separately in each z-slice and then average over z-slices in order to accurately determine the nuclear to cytoplasmic ratio. For all other experiments, calculations were performed on maximum intensity projections.

### Cell Tracking

For tracking experiments, MIXL1-expressing cells were identified using Ilastik’s pixel classification tool to generate both probability and segmentation masks. Tracking was performed using the Tracking-Pixel prediction map in Ilastik, and results were exported as CSV files. These data were subsequently analyzed using custom MATLAB scripts.

### Bulk RNA Sequencing

hESCs were seeded at a density of 30,000 cells/cm² in experimental and control conditions. Control samples were maintained in mTeSR1 supplemented with 10 μM RI and other samples were grown in AMLC differentiation media described above for 2 or 4 days. Cells were harvested as single-cell suspensions using Accutase, and total RNA was extracted the same day using the RNaqueous kit (Invitrogen). RNA quality was assessed using agarose gel electrophoresis and an Agilent 2000 system to confirm integrity and the absence of DNA contamination. Sequencing was performed on the Illumina NovaSeq6000 platform with a target depth of 6 Gb per sample.

### Bulk RNA sequencing data processing

To quantify the abundance of transcripts for each sample, Salmon v1.9.01 was used. Reads were aligned to the human transcriptome (GRCh38) index for salmon, with salmon index using the selective alignment method (salmon_sa_index:default in http://refgenomes.databio.org/v3/genomes/splash/2230c535660fb4774114bfa966a62f823fdb6d21acf138d4). We quantified transcripts with Salmon using the −l A flag to infer the library type (paired-end) automatically and the –validateMappings flag to use the selective alignment method. The resulting datasets were then processed using the tximeta2 package in R, and the counts matrix was exported for downstream analyses.

### Comparisons to published single-cell RNA sequencing datasets

Bulk RNA sequencing data was compared to single-cell RNA sequencing datasets published in Oldak et al. 2023 and Yang et al. 2021. All reference data used were publicly available with published cell type annotations. Cynomolgus monkey ENSEMBL ids in Yang et al. 2021 were converted to human gene symbols using biomaRt. Reference datasets were first pre-processed and analyzed as described in Oldak et al. 2023 and Yang et al. 2021, respectively. Then, to relate bulk samples from our in vitro sequencing data to the Oldak et al. 2023 and Yang et al. 2021 databases, we projected our bulk RNA sequencing data onto the single-cell RNA sequencing datasets using latent semantic indexing projection (lsi) as proposed in ^27^. We also compared aggregate and averaged expression for a subset of marker genes of cell types of interest identified in the single-cell RNA sequencing datasets to the mean expression of the bulk RNA sequencing data samples.

### Data sources

Previously published data are publicly available. Data were from a human *in vitro* stem cell based embryo model^26^ (GEO accession GSE239932), and from cynomolgus monkey^10^ (GEO accession GSE148683).

## Acknowledgements

We thank Ed Stanley (Murdoch Childrens Research Institute, Australia) for the MIXL1 cell line and members of the Warmflash lab for helpful discussions. This work was supported by grants from NSF (MCB-2135296) and NIH (R01HD112488) to AW.

