## Supplementary Figures and Table for "Juxtaposition of human pluripotent stem cells with amnion-like cells is sufficient to trigger primitive streak formation"

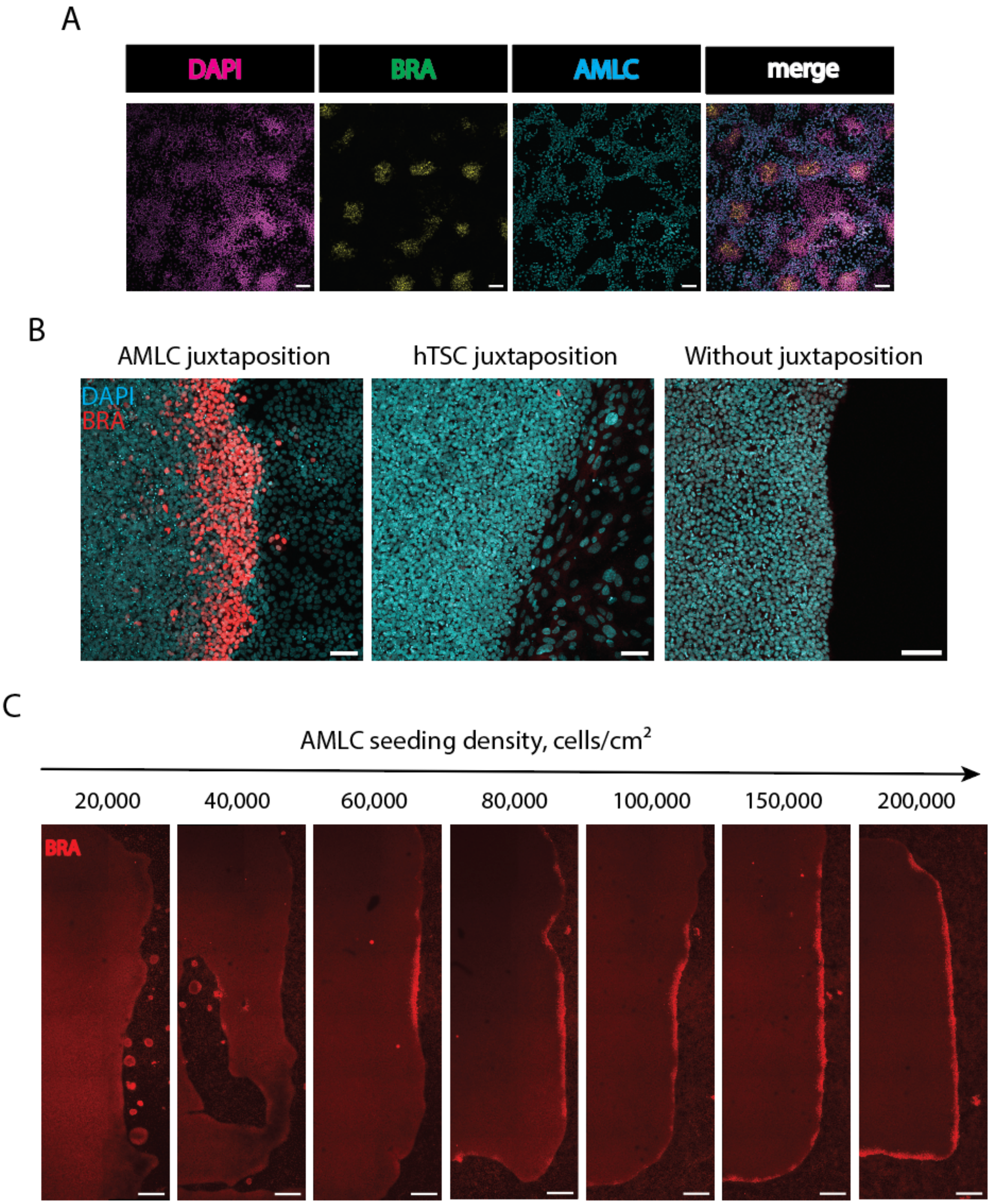


**Figure S1. Induction is a property of AMLCs, but not hTSCs, and requires high cell density.** (A) Representative image of an experiment mixing AMLCs (labeled by H2B-Cerulean) and hPSCs, culturing for 48 hours, and immunostaining for BRA. Scale bar, 100μm. (B) Representative immunostaining for BRA in an AMLC-hPSC juxtaposition compared to a hTSC-hPSC juxtaposition, and one in which no cells are juxtaposed with the hPSCs. hTSCs are seeded at the same density as the AMLCs. Scale bar, 100 μm. (C) Stitched images of a large area of juxtaposed AMLCs and hPSCs showing immunostaining of BRA. AMLCs are seeded with the indicated density and fixed 60 hours after juxtaposition. A portion of the 200,000 seeding density group is shown in figure 1D. Scale bar, 500 μm

A

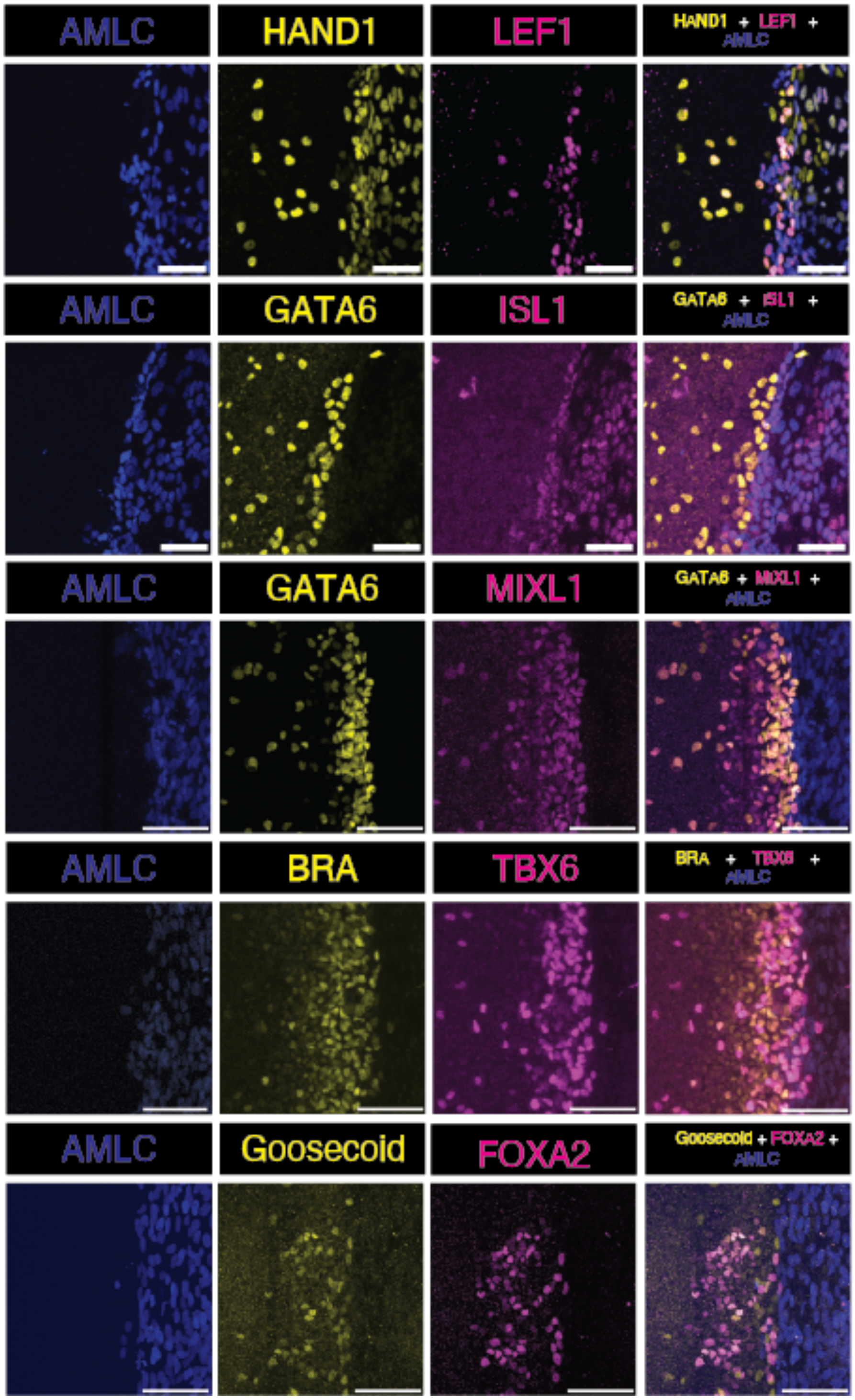


**Figure S2. AMLCs induce the differentiation of a wide array of mesendodermal cell types.** Representative immunostaining of juxtaposition experiments for the indicated markers at the end of day 4. Scale bars: 100µm.

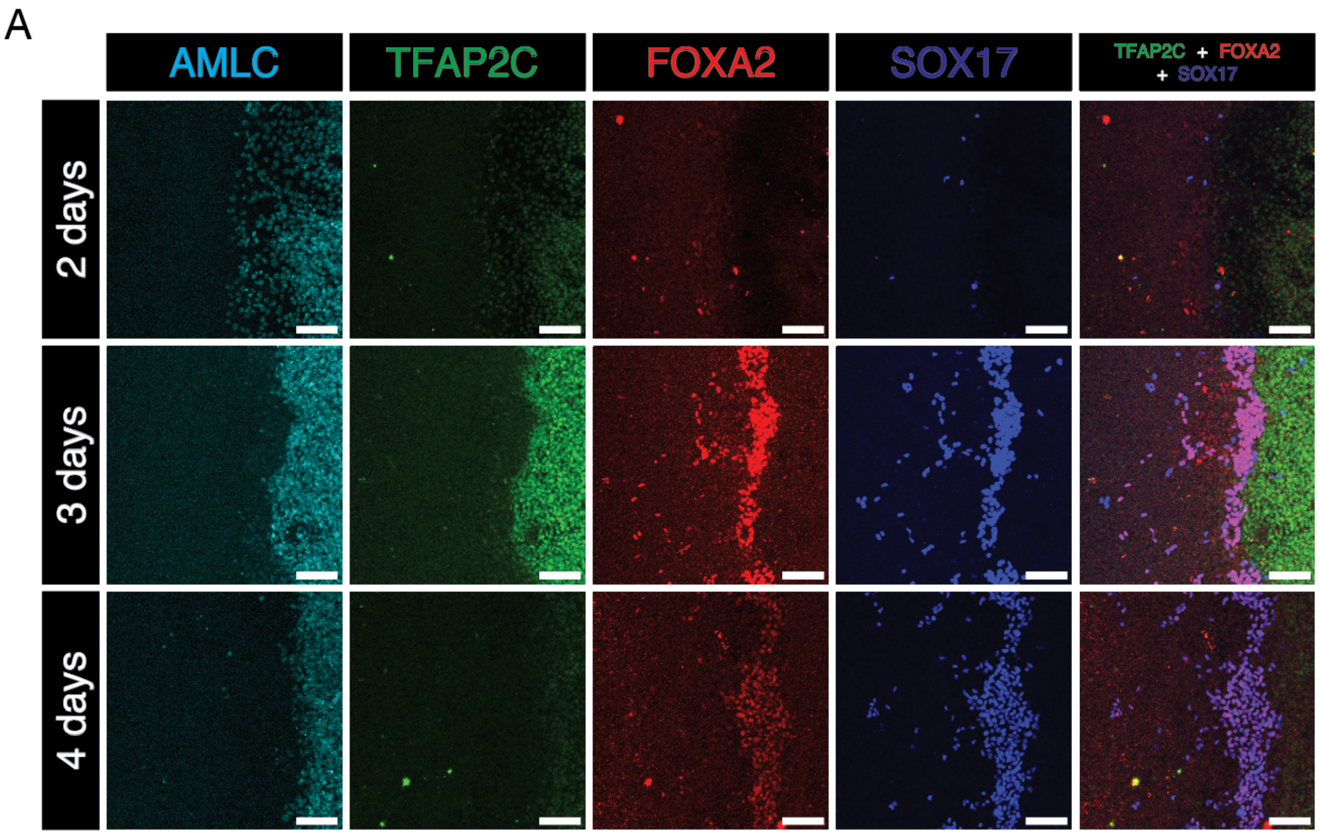


**Figure S3. Dynamics of endoderm differentiation in juxtaposition experiments**. Representative immunostaining for the indicated markers in the juxtaposition experiment at the end of day 2, 3, or 4. Scale bars: 100μm.

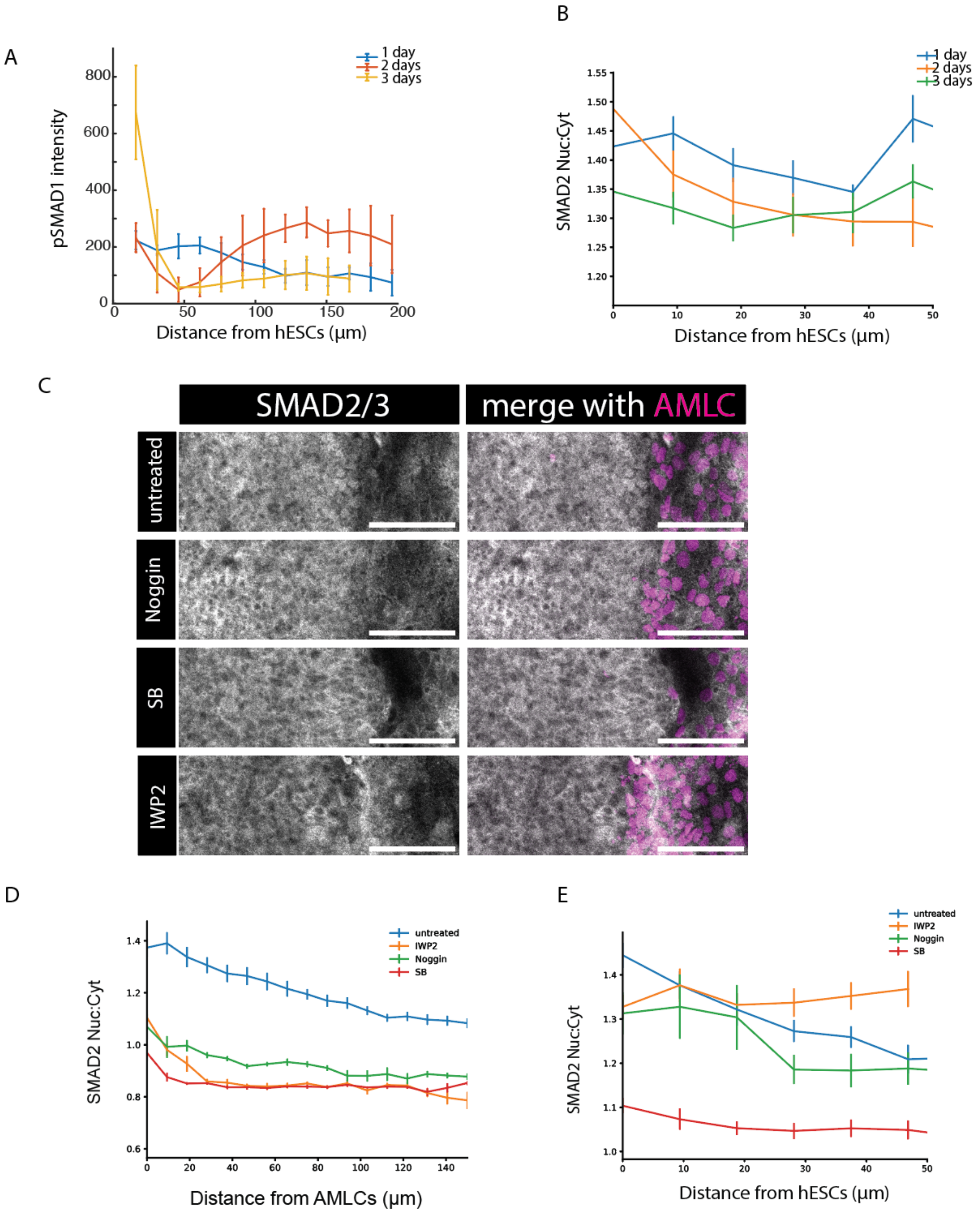


**Figure S4. Quantification of SMAD signaling in the AMLC compartment.** (A, B) Quantification of signaling in the AMLCs in the experiments shown in Figure 4A and 4C. pSMAD1 intensity (A) or SMAD2 nuclear to cytoplasmic ratio (B) is quantified in the AMLCs and plotted as a function of the distance from the hPSCs. (C-E) Representative immunostaining image (C) and quantification of the nuclear to cytoplasmic ratio in the hPSC (D) and amnion (E) compartments of SMAD2/3 after 2 days of juxtaposition with BMP, Nodal, or Wnt signaling inhibited by Noggin, SB or IWP2, respectively. Error bars: s.e.m. n>5. Scale bars: 100 µm.

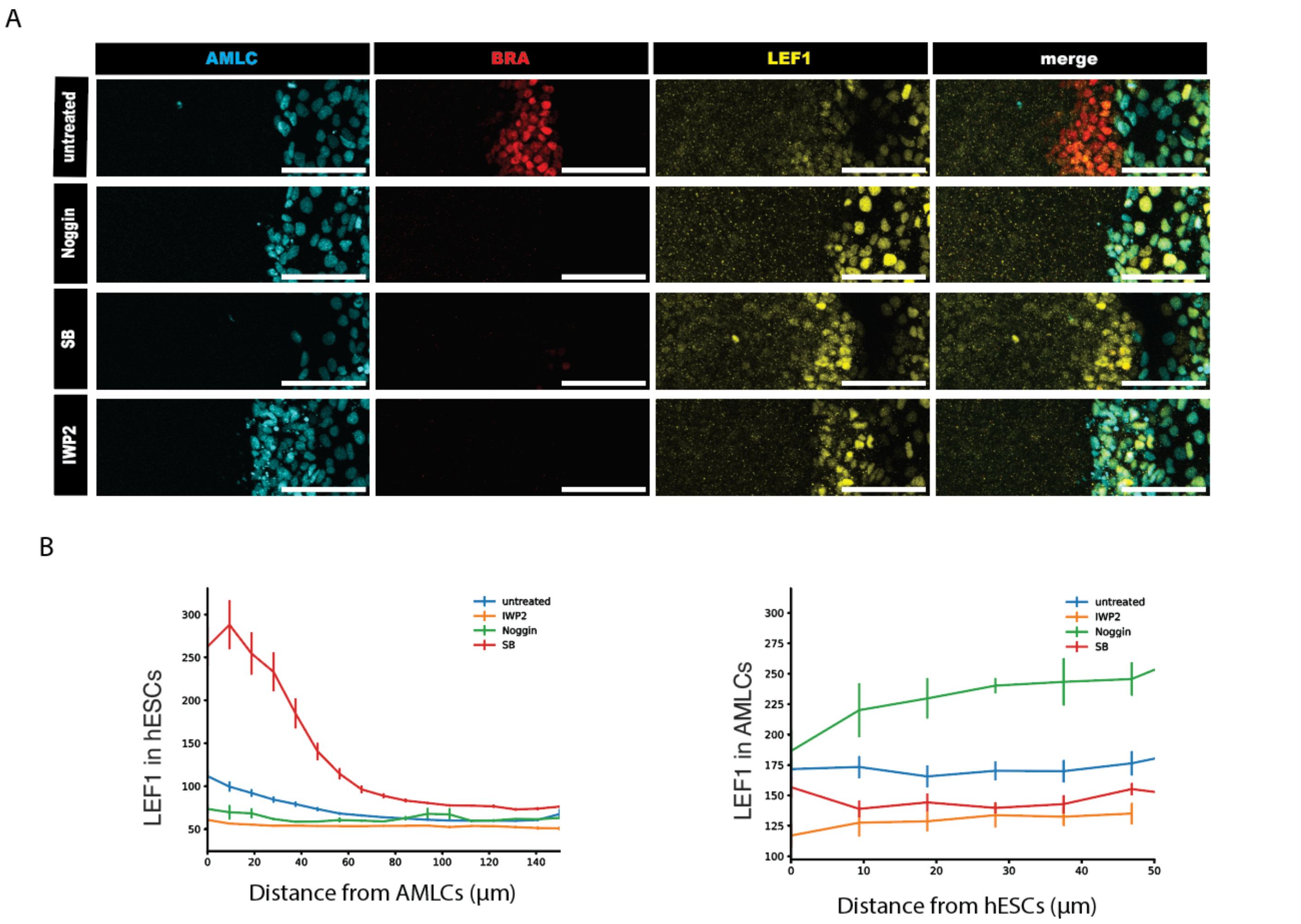


**Figure S5. Effect of pathway inhibition on LEF1 expression.** (A, B ) Representative immunostaining (A) and quantification (B) of LEF1 immunofluorescence after 1, 2, or 3 days of juxtaposition with pathway inhibitors as indicated. Scale bars: 100 μm. Error bars: s.e.m. n > 5.

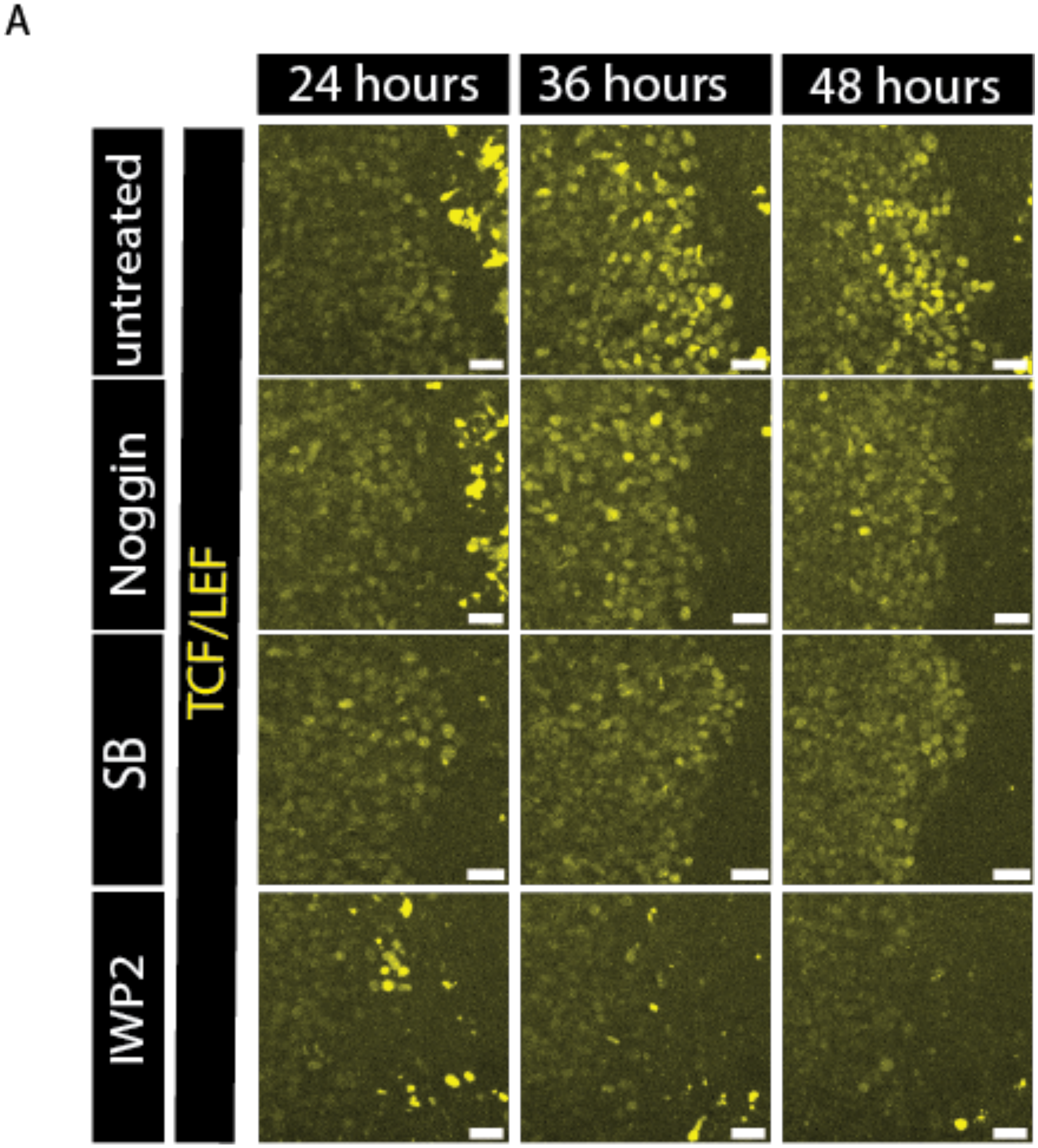


**Figure S6. Time lapse imaging of WNT dynamics with pathway inhibitors.**
Snapshots of the juxtaposition experiment where TCF/LEF::GFP reporter cells were used in the hPSC compartment at the indicated time points during live imaging.

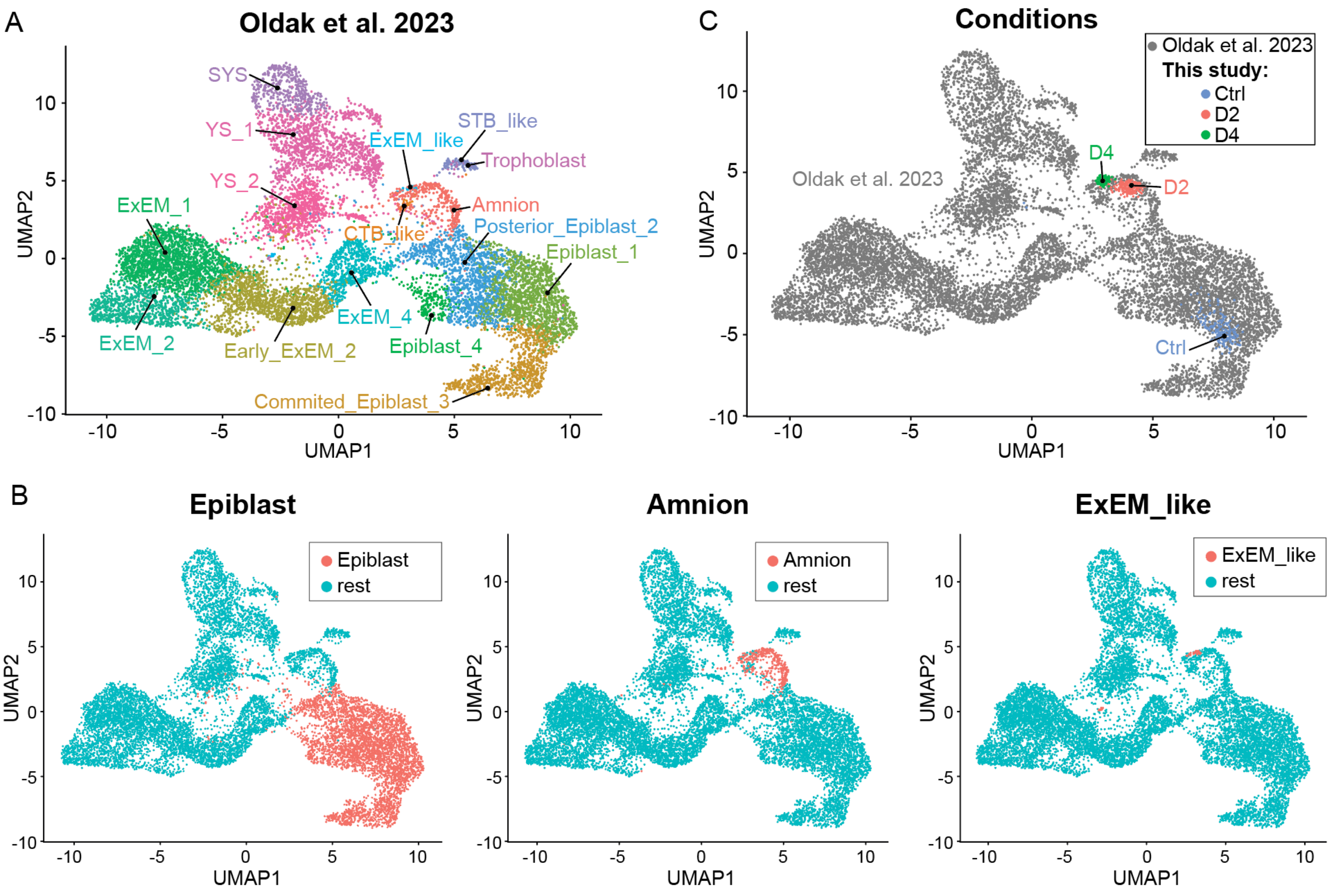


**Figure S7. Comparing AMLC differentiation with cell types in a complete human embryo model (Oldak et al., 2023).** (A) (left) UMAP of data from Oldak et al with cell types indicated. (right) Data from this study at 0, 2, or 4 days of differentiation mapped to the Oldak et al dataset (gray points). (B) Highlighting epiblast, amnion, and ExEM-like cells in the Oldak data.

| Antibody Name | Manufacturer | Catalog Number | Dilution | Species |
|---|---|---|---|---|
| Alexa Fluor 488 anti-Goat | Life Technologies | A11055 | 1:500 | Donkey |
| Alexa Fluor 488 anti-Mouse | Life Technologies | A21202 | 1:500 | Donkey |
| Alexa Fluor 488 anti-Rabbit | Life Technologies | A21206 | 1:500 | Donkey |
| Alexa Fluor 555 anti-Goat | Life Technologies | A21432 | 1:500 | Donkey |
| Alexa Fluor 555 anti-Mouse | Life Technologies | A31570 | 1:500 | Donkey |
| Alexa Fluor 555 anti-Rabbit | Life Technologies | A31572 | 1:500 | Donkey |
| Alexa Flour 647 anti-Goat | Life Technologies | A21447 | 1:500 | Donkey |
| Alexa Fluor 647 anti-Mouse | Life Technologies | A31571 | 1:500 | Donkey |
| Alexa Fluor 647 anti-Rabbit | Life Technologies | A31573 | 1:500 | Donkey |
| BRACHYURY | R&D Systems | AF2085 | 1:300 | Goat |
| GATA-6 | R&D Systems | AF1700 | 1:300 | Goat |
| GOOSECOID | R&D Systems | AF4086 | 1:100 | Goat |
| HAND1 | R&D Systems | AF3168 | 1:200 | Goat |
| SOX17 | R&D Systems | AF1924 | 1:200 | Goat |
| TBX6 | R&D Systems | AF4744 | 1:200 | Goat |
| GFP | Abcam | ab1218 | 1:200 | Mouse |
| ISL1 | DSHB | 3B5 | 1:100 | Mouse |
| SMAD2/3 | BD Biosciences | 610843 | 1:200 | Mouse |
| BRACHYURY | R&D Systems | MAB20851 | 1:400 | Rabbit |
| CDX2 | Cell Signaling Technology | 12306s | 1:100 | Rabbit |
| FOXA2 | Abcam | ab108422 | 1:300 | Rabbit |
| LEF1 | Cell Signaling Technology | 2230s | 1:200 | Rabbit |
| pSMAD1/5/8 | Cell Signaling Technology | 13820 | 1:400 | Rabbit |
| SMAD2/3 | Cell Signaling Technology | 5339S | 1:200 | Rabbit |
| SOX2 | Cell Signaling Technology | 23064s | 1:200 | Rabbit |

**Supplementary Table 1. Antibodies used in this study.**
